# Maize immune signalling peptide ZIP1 evolved de novo from a retrotransposon

**DOI:** 10.1101/2022.05.18.492421

**Authors:** Jasper R.L. Depotter, Johana C. Misas Villamil, Gunther Doehlemann

## Abstract

Plants are subjected to different types of threats that require appropriate physiological responses to counteract them. Signalling peptides are produced under specific conditions and elicit physiological changes. *PROZIP1* encodes such a signalling peptide, Zip1, that induces salicylic acid defence responses in maize (*Zea mays*) leading to a better protection against biotrophic pathogens. Despite salicylic acid pathway being conserved amongst plants, we observed that Zip1 only occurs in the *Zea* genus. *PROZIP1*’s evolution is associated with transposons, as it resides in the terminal repeat of a retrotransposon from the Gyma family. We traced back the mutations that were encountered by this transposon and found that *PROZIP1* emerged *de novo* in *Zea*. This emergence likely occurred less than 728,000 years ago. In conclusion, we describe the evolution of a recently emerged plant immune signalling peptide from a transposon sequence.

## Introduction

Transposable elements (TEs) are mobile DNA fragments that occur in genomes of virtually all walks of life, even in giant viruses (1). They were first identified in maize following the observation of differences in anthocyanin pigment formation and were initially called “controlling elements” (2). Although their importance was initially marginalized as “junk DNA”, TEs are currently acknowledged as important drivers of genome evolution (3). The ability of TEs to move within genomes creates variation that can be detrimental, but also provide opportunities to adapt to new environmental conditions.

Transposable elements come in many different forms and shapes (4). There are two main classes of transposable elements, distinguished on the transposition intermediate being RNA (class I) or not (class II). These two classes have also been referred to as retrotransposons and DNA transposons, respectively. TEs are then further classified based on their structure (5). One type of retrotransposons is long terminal repeat retrotransposon (LTR-RT) that is characterized by a direct repeat at each end of their sequence (6). LTR-RTs produce a polycistronic transcript for *gag, pol* and, in case of retroviruses, *env*. The *gag* gene encodes for a structural protein to form the capsid where the reverse transcription occurs. The *pol* gene encodes for a protease, a reverse transcriptase and an integrase. Based on the order of protein domains in the pol protein and sequence similarity LTR-RTs can be further distinguished in the superfamilies Ty1-*Copia* and Ty2-*Gypsy*. If the enzymatic machinery of gag and pol proteins is encoded, LTR-RTs can be autonomous for their transposition. Nonautonomous LTR-RTs, without this enzymatic machinery, can still be mobile but depend and the enzymatic machinery of other TEs.

The genome fraction that consists of TEs can differ significantly between plant species. Less than 20% of the *Arabidopsis thaliana* genome consists of TEs, whereas this is more than 80% for the maize genome (7–10). Also, within plant species, substantial variation in TEs can be observed through recent transpositions as well as deletions of TEs (11). In four maize lines, 400,000 of such polymorphic TEs were found, of which only 111,000 elements were found in all four lines (12). Partly due to the first TE discovery, maize has been a model system for TE research. Generally, almost ninety percent of TEs in maize belong to the class I of retrotransposons (7, 10). Around one third of the classified retrotransposons are of the Ty1-*Copia* variant and the other two thirds of the *Gypsy* variant. Unsurprisingly, TEs played a pivotal role in the evolution and domestication of maize (13). Maize was domesticated in southern Mexico about 9,000 years ago with teosinte *Zea mays* ssp. *parviglumis* being the closest wild progenitor that diverged ∼55,000 years ago from maize (14, 15). After domestication, maize has regained diversity through introgression from wild teosinte *Zea mays* spp. *mexicana* that diverged from maize ∼60,000 years ago (14, 16, 17).

To prevent uncontrolled transposition, hosts keep the expression of TEs in check through epigenetic means (18). However, TE transpositions can also be a source of genomic variation, which can sporadically lead to environmental adaption. TE sequences themselves are also subjected to evolution and may evolve a function that is advantageous for its host (domestication). This is illustrated by a LTR-RT in one of the introns of the *Arabidopsis* gene *RPP7*, which confers resistance to the downy mildew pathogen *Hyaloperonsopora arabidopsis* (19). The extent to which this LTR-RT is silenced impacts the choice of two alternative *RPP7* polyadenylation sites leading to different transcripts. One transcript encodes RPP7 and one does not, which gives *Arabidopsis* the opportunity to finetune *RPP7* expression in response to pathogen attack. Hosts can also domesticate TEs into the production of coding and non-coding RNAs that provide evolutionary advantages (4, 20). TEs can be domesticated into the production of microRNAs that act as part of an immune response against pathogen invasion. In wheat, microRNAs with induced expression upon powdery mildew (*Blumeria graminis* f.sp *tritici*) infection frequently resided in miniature inverted-repeat TEs of the *Mariner* superfamily (21). Alternatively, TE proteins can be domesticated into proteins that play a role in host immune systems (22). In the adaptive immune system of jawed vertebrate, a variety of antibodies and T cell receptors are produced by RAG1-RAG2 recombinase and V(D)J recombination, which evolved from a DNA transposon (23). The RAG1-RAG2 recombinase evolved from the transposase gene of the ancestor transposon (24). Furthermore, TE domestication seems also to have played a pivotal role in the evolution of the CRISPR-Cas adaptive immune system of prokaryotes, as Cas1 endonuclease originates from casposons, a superfamily of archeal and bacterial transposons (25).

Plants have a plethora of peptides that activate stress responses upon detection of abiotic and biotic stresses (26). *Zea mays* immune signalling peptide 1 (Zip1) is a phytocytokine that elicits salicylic acid (SA)-dependent defence signalling in maize (27). SA signalling promotes efficient defence activation against biotrophic pathogens but may facilitate the colonization of necrotrophic pathogens that benefit from cell death (28).

Indeed, Zip1 promotes the infection of the necrotrophic fungus *Botrytis cinerea* and reduces virulence of the biotrophic fungus *Ustilago maydis* (27). Zip1 is derived from the precursor protein PROZIP1. Intriguingly, despite the activation of a conserved immune signalling pathway in plants, *PROZIP1* is an orphan gene as we did not find significant homology in the NCBI database, neither with nucleotide sequence nor with protein sequence blast (consulted 23/02/2022). This triggered our interest to investigate how *PROZIP1* emerged.

## Results

### *PROZIP1* occurs at different genome locations

To know the genomic environment of *PROZIP1*, we looked where in the maize genome this gene is located. Recently, the 25 founder lines of the Maize Nested Association Mapping (NAM) population were sequence and assembled, in addition to a new version of the maize reference line B73 (10). Two of the 26 screened maize lines, P39 and Ky21, did not contain an open reading frame in their genome assembly that could encode PROZIP1 (Table 1). In the other maize lines, *PROZIP1* occurs in one (7 lines), two (10 lines) or three copies (7 lines). Except from one copy in maize line NC358 (R65Q, 194G>A), all PROZIP1 copies have the identical protein sequence. On nucleotide levels, there are two more alleles with synonymous substitutions: 87G>A in two maize lines (Mo18W and CML247) and 120C>T in three maize lines (CML69, M37W and CML322). *PROZIP1* occurs also as a pseudogene in nine maize lines with three different alleles (Table 1) due to indels (insertions or deletions) that cause premature stop codons (Figure S1). *PROZIP1* genes and pseudogenes can occur on different chromosomes, as copies are found on chromosome 7, 8, 9 and 10. On chromosome 10, *PROZIP1* (pseudo)genes always occur in two copies that reside 8.3 kb or 20.3 kb from each other, whereas on chromosome seven *PROZIP1* always occurs as a single gene (Table 1, Figure S1). *PROZIP1* was only found in NAM line Oh7B on chromosome 9, similar to chromosome 10, with two copies that have 8.3 kb between each other (Table 1, Figure S1). Only *PROZIP1* pseudogenes were found on chromosome eight in one or two copies (Table 1, Figure S1). These two copies are 7.8 kb apart from each other. In conclusion, the different copy numbers of *PROZIP1* in different maize lines on various genome location indicate that *PROZIP1* had a dynamic evolution with duplications and translocations.

**Table 1.** Chromosome location of *PROZIP1* genes (G) and pseudogenes (P) in maize B73 and NAM lines. The NAM lines are coloured based on their primary grouping (gold = stiff stalk heterotic group, blue = non-stiff-stalk heterotic group, grey = mixed tropical-temperate ancestry, purple = popcorn, orange = sweet corn, green = tropical)

| Maize line | Chr7 | Chr8 | Chr9 | Chr10 |  |
| --- | --- | --- | --- | --- | --- |
|  | G | P | G | G | P |
| B73 | 1 | 2 |  |  |  |
| B97 | 1 | 2 |  | 2 |  |
| Ky21 |  |  |  |  |  |
| M162W | 1 |  |  |  |  |
| Ms71 |  |  |  | 2 |  |
| Oh43 |  |  |  | 2 |  |
| Oh7B |  |  | 2 |  |  |
| M37W | 1 | 2 |  |  |  |
| Mo18W | 1 |  |  |  |  |
| Tx303 | 1 |  |  | 2 |  |
| HP301 |  |  |  | 2 |  |
| II14H |  |  |  | 2 |  |
| P39 |  | 1 |  |  |  |
| CML103 |  |  |  | 2 |  |
| CML228 | 1 |  |  | 2 |  |
| CML247 | 1 |  |  | 1 | 1 |
| CML277 |  |  |  | 1 | 1 |
| CML322 | 1 |  |  | 1 | 1 |
| CML333 | 1 |  |  | 2 |  |
| CML52 | 1 |  |  | 1 | 1 |
| CML69 | 1 |  |  | 2 |  |
| Ki11 | 1 |  |  | 2 |  |
| Ki3 | 1 |  |  |  |  |
| NC350 | 1 |  |  | 2 |  |
| NC358 |  |  |  | 2 |  |
| Tzi8 |  |  |  | 1 | 1 |

\**PROZIP1* (pseudo)genes on chromosome 10 that reside 20.3 kb from each other.

### *PROZIP1* resides in the terminal repeat of an LTR-RT

As *PROZIP1* occurs in different locations of the maize genome, we examined if this gene is associated with transposable elements. Using the maize transposable element annotations by Hufford et al. (2021), we found that *PROZIP1* on chromosome 10 is part of the LTRs of an LTR-RT from the Gypsy LTR-RT family Gyma (Figure 1A). In contrast, *PROZIP1* on chromosome 7 resides inside an LTR-RT from the Ty1-*Copia* LTR-RT family Opie. The *PROZIP1 (pseudo-)*gene locations on chromosome 8 and 9 were not annotated as a TE by Hufford et al. (2021). However, *PROZIP1* LTR-RT on chromosome 10 displays high homology to the genome region where *PROZIP1 (pseudo-)*genes are located on chromosome 8 and 9 (Figure 1B). Consequently, *PROZIP1 (pseudo-)*genes on chromosome 8 and 9 are also part of the LTRs of an unannotated LTR-RT, which is homologous to the one on chromosome 10.

**Figure 1.**
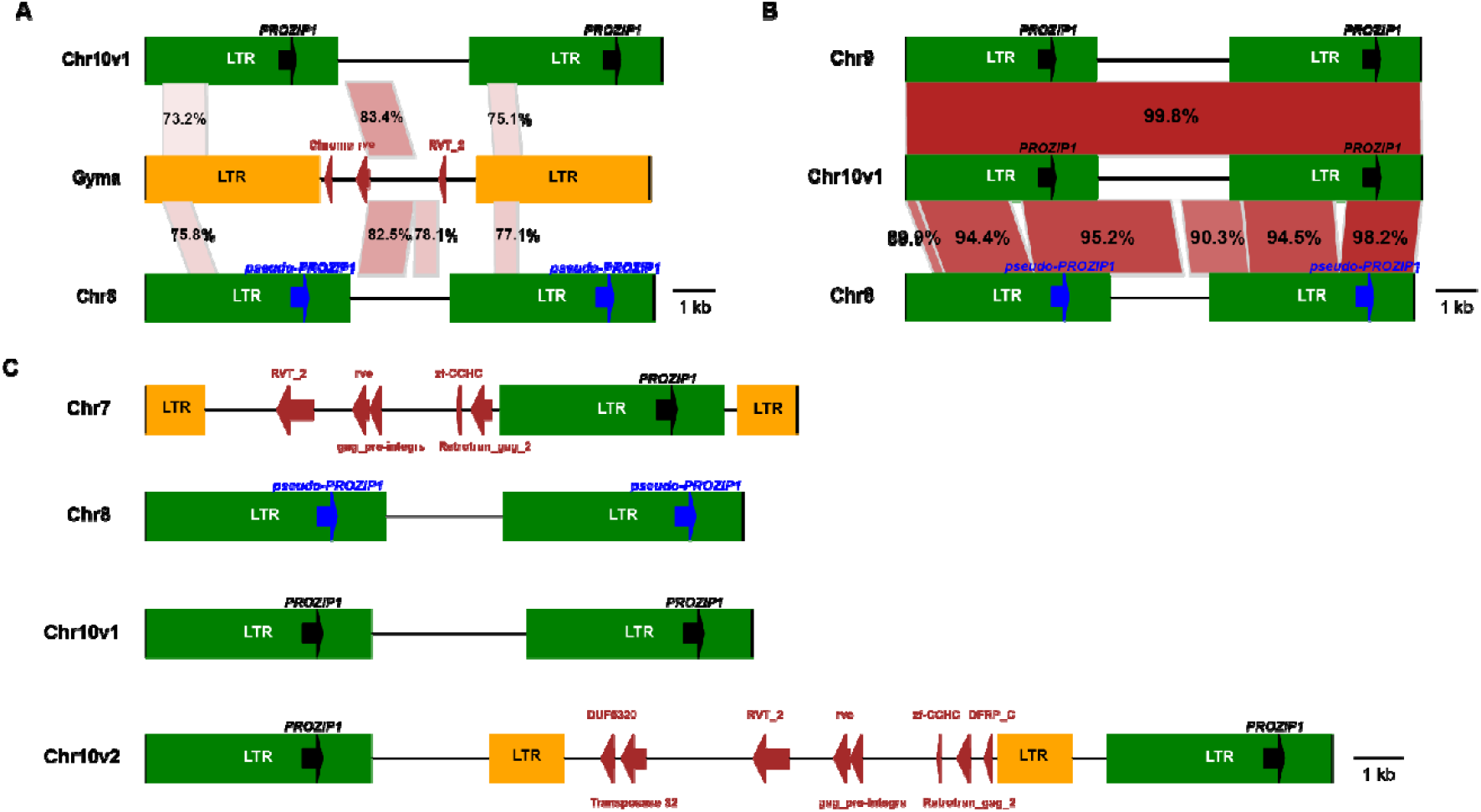
*PROZIP1* flanking regions at different genome locations. Green box indicates the long terminal repeat (LTR) where the *PROZIP1* (pseudo)gene resides. Yellow box indicates the LTR region of other retrotransposons. Red arrow indicates the location and orientation of a functional domain annotated in an open reading frame. (A) Sequence identity between *PROZIP1* LTR-RTs and the representative of the Ty2-*Gypsy* LTR-RT *Zea mays* Gyma family from the TREP database (57). (B) Homology between the *PROZIP1* LTR-RT alleles on chromosome 8, 9 and 10. (C) The different LTRs where *PROZIP1* resides. Sequence identities were obtained using BLASTN (2.2.31+).

The LTR where *PROZIP1* resides has a size of 4.6 kb for the chromosome 9 and 10 alleles and 4.9 kb on chromosome 8. We could not identify open reading frames with functional domains in the regions between these LTRs (Figure 1C). On chromosome 7, *PROZIP1* is flanked by the same 4.6 kb LTR sequence, which is embedded in a Ty1-*Copia* LTR-RT (Figure 1C). As previously mentioned, distances between (pseudo-)*PROZIP1* copies on chromosome 10 can be 8.3 kb or 20.3 kb. In the 20.3 kb variant, a Ty1-*Copia* LTR-RT inserted in the *PROZIP1* LTR-RT (Figure 1C). Across B73 and the NAM lines, the LTR sequence flanking *PROZIP1* displays 56 variants (SNPs or indels) per 1kb, whereas this is 7 per 1 kb for the *PROZIP1* coding regions (Figure S2). Thus, *PROZIP1* is subjected to a higher degree of conservation than its flanking sequences, which indicates its functional importance for maize.

### *PROZIP1* emerged as a new gene

As we could not find any *PROZIP1* ortholog in the NCBI database, we investigated the emergence of *PROZIP1* as a signalling peptide encoding gene. We identified homologous LTR-RT to the *PROZIP1* LTR-RT using the maize TE annotation of maize NAM and B73 lines (10). We selected LTR-RTs that have at least 70% sequence identity in 55% of the sequence reciprocally with the *PROZIP1* LTR-RT allele on chromosome 10, which were 407 LTR-RTs in total. As the chromosome 8 allele of *PROZIP1* LTR-RT, is not annotated in the TE annotation of B73, B97 and M37W, we added these three alleles to our list. Consequently, we removed identical LTR-RTs (minimum sequence identity and overlap of 99% and 80%, respectively), which resulted in a final list of 97 LTR-RTs (Figure 2A). Using these LTR-RT homologs, we reconstructed the emergence of *PROZIP1* as part of an LTR-RT. Based on Maximum Likelihood, all the sequences of the *PROZIP1* LTR-RT ancestors were reconstructed. We selected the *PROZIP1* LTR-RT ancestors and consequently aligned them to see if *PROZIP1* evolved through gradual mutagenesis or if the complete *PROZIP1* gene was recently inserted in the LTR-RT terminal repeat region at once. The alignment of the ancestor sequences to (*pseudo-*)*PROZIP1* LTR-RT sequences suggests that the *PROZIP1* gene was not recently inserted. Ancestor sequences do not display a complete absence of *PROZIP1* but rather *PROZIP1* gradually evolved through numerous indels and nucleotide substitutions (Figure 2B). In conclusion, *PROZIP1* likely evolved *de novo* through sequential nucleotide mutations in the terminal repeat region of an LTR-RT.

**Figure 2.**
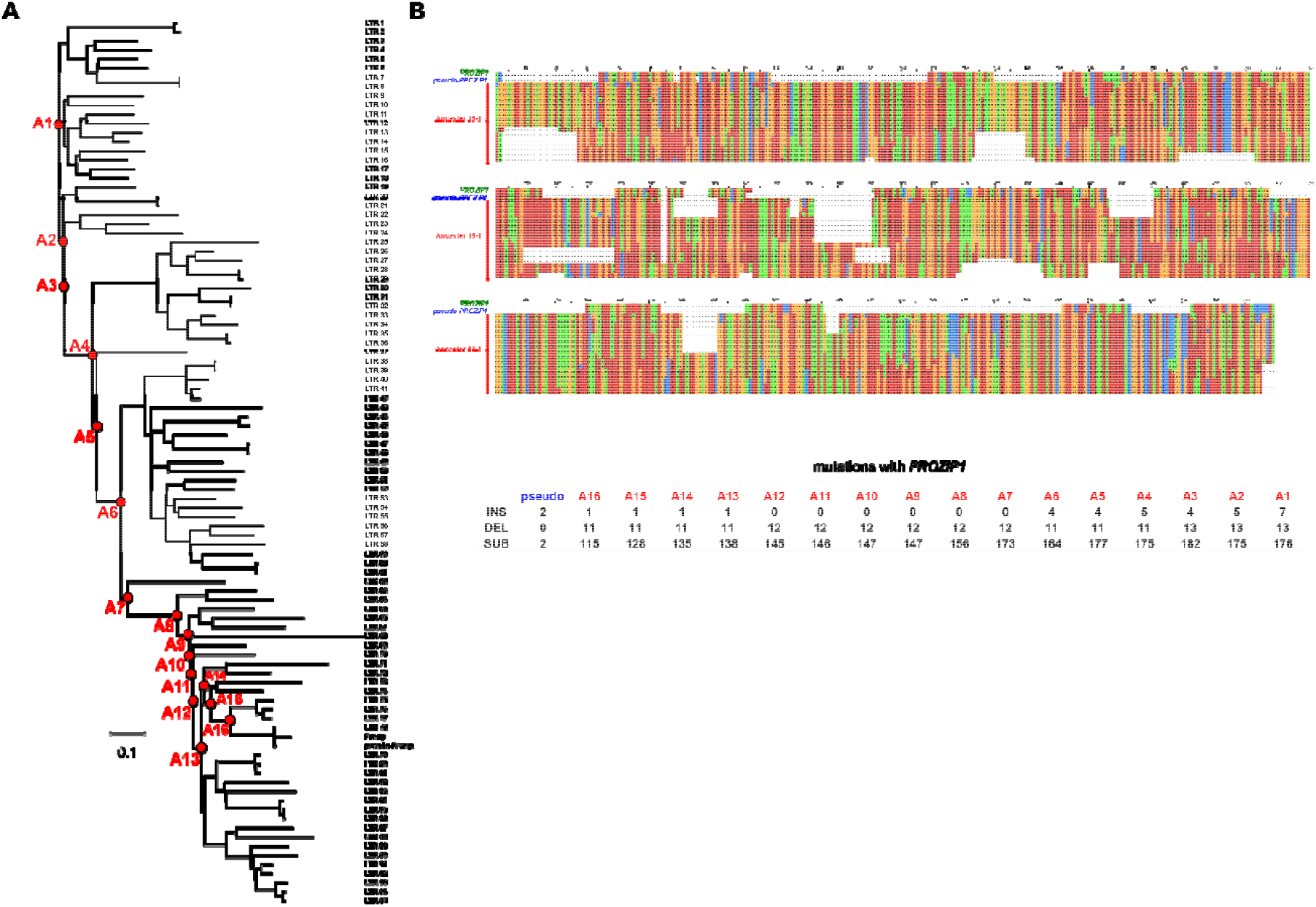
The emergence of *PROZIP1* in the terminal repeat of an LTR-RT. (A) Phylogenetic tree of unique LTR-RTs that have at least 70% sequence identity in 55% of the sequence reciprocally with the *PROZIP1* LTR-RT. All ancestors that are aligned in section B are indicated by a red dot. (B) Ancestor sequences were aligned with the (*pseudo-*)*PROZIP1* LTR-RTs of which the sequence corresponding to *PROZIP1* is displayed. INS, DEL and SUB represent the number of insertions, deletions and substitutions of the ancestors and *pseudo-PROZIP1* with *PROZIP1*.

### PROZIP1 encodes a *Zea* specific signalling peptide

As gene annotations of some closely related (sub-)species of *Zea mays* are not available, we screened genome sequences and mapped reads of these close relatives to see if *PROZIP1* is present. *PROZIP1* is present in a single copy in a publicly available genome assembly of teosinte *Zea mays* subsp. *mexicana* (Accession: PRJNA299874) (29). The nucleotide sequence of *PROZIP1* in *Zea mays* subsp. *mexicana* is 100% identical to that of the most prevalent *PROZIP1* in *Zea mays* subsp. *mays.* We also mapped publicly available reads from a whole genome sequencing project (Accession: PRJNA300309) of the closest progenitor of cultivated maize, teosinte *Zea mays* subsp. *parviglumis*, to the genome assembly of maize line Ky21 (lacks *PROZIP1*, Table 1) with the addition of the *PROZIP1* LTR-RT terminal repeat that contains *PROZIP1* (10). Reads of *Zea mays* subsp. *parviglumis* were mapped and completely covered *PROZIP1*, indicating the presence of *PROZIP1* in the closest progenitor of cultivated maize. Eastern gamagrass, *Tripsacum dactyloides*, is a close relative of *Zea mays*, of which several whole-genome sequencing projects have been performed (Accessions: SRP011907, SRP095374, SRP158906) (30). In none of the *Tripsacum dactyloides* accessions reads were mapped to *PROZIP1*. Thus, *PROZIP1* is absent in the hitherto sequenced *Tripsacum dactyloides* genomes. These data indicate, that *PROZIP1* emerged before maize domestication as teosinte progenitors of maize contain *PROZIP1*, but no orthologs in other plant species have been found.

To determine more precisely when *PROZIP1* emerged, we want to date the nodes of the phylogenetic tree of the different *PROZIP1* variants. We use the LTR region containing *PROZIP1* to construct the tree, as this region is present in all genome locations. To include the closest wild progenitor *Zea mays* subsp. *parviglumis,* the variants of the mapped *Zea mays* subsp. *parviglumis* reads to *PROZIP1* LTR-RT terminal repeat were called and a new consensus sequence was constructed. Two different haplotypes were constructed that are 99.7% identical and both contain *PROZIP1* that is 100% identical to *PROZIP1* of maize. To root the tree, LTR-RT Chrom10:23797712-23810820 from B97 was used, which is a close homolog of (*pseudo*-)*PROZIP1* LTR-RT that contains the LTR where *PROZIP1* resides. The teosinte alleles from the subspecies mexicana and parviglumis do not form an outgroup, rather they cluster together with the *PROZIP1* alleles on chromosome locations 9 and 10 (Figure 3). Moreover, despite that subspecies mexicana diverged a longer time ago from maize than parviglumis (14), the mexicana allele seemed to have diverged more recently from the cultivated maize allele than parviglumis. The allele on chromosome 7 diverged more recently from the chromosome 10 allele than the chromosome 8 allele.

**Figure 3.**
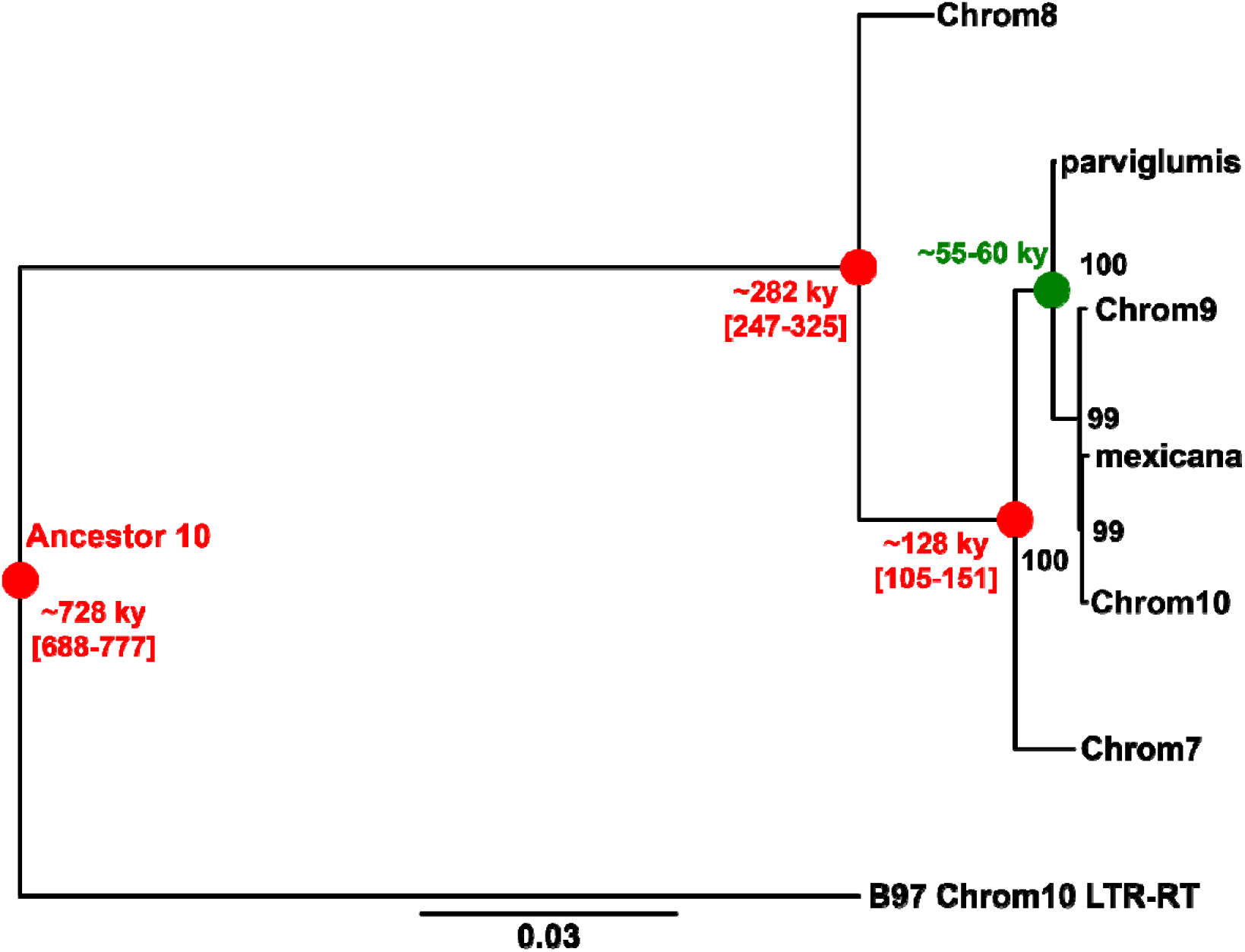
Phylogenetic relationship and node dating of the different (pseudo-)*PROZIP1* alleles. The terminal repeats wherein (*pseudo*-)*PROZIP1* resides were used for tree construction. Nodes were dated by using the last common ancestor of Chrom10 and parviglumis as a calibration node (green). The mean age of the nodes is reported together with the 95% credibility interval. Parviglumis and mexicana represent the allele of *Zea mays* ssp. *parviglumis* and *Zea mays* ssp. *mexicana*, respectively. The robustness of the inferred phylogeny was assessed by 100 bootstrap approximations. B97 LTR-RT on chromosome 10 located between 23797712 and 23810820 was used as an outgroup to root the tree.

To estimate the timeframe at which *PROZIP1* emerged, we dated the nodes of the phylogenetic tree. Here, the last common ancestor of the chromosome 10 and parviglumis alleles was used as a calibration node. Subspecies *parviglumis* and *mexicana* diverged around 55,000 and 60,000 years ago, respectively (14). As we cannot exclude that the chromosome 10 allele originates from subspecies *mexicana*, of which genome regions were introgressed during maize domestication around 9,000 years ago (15, 31), we performed the node dating with the calibration point as a uniform distribution between 55,000 and 60,000 (Figure 3). The last common ancestor of all the *PROZIP1* alleles is calculated ∼128,000 years ago. *Pseudo-PROZIP1* LTR-RT and the *PROZIP1* LTR-RT diverged ∼282,000 years ago, whereas divergence was estimated to be ∼728,000 years ago for *PROZIP1* LTR-RT and the B97 outgroup LTR-RT indicating that *PROZIP1* evolved less than 728,000 years ago.

## Discussion

TEs play a pivotal role in the adaptive potential of many eukaryotes to different stress factors (32). *PROZIP1* encodes a maize immune signalling peptide that likely evolved within an LTR-RT through various mutations (Figure 3). Its evolution within a transposon has very likely facilitated *PROZIP1* mobility within the maize genome, through transposition or through ectopic recombination of homologous sequences (33). Hence, it may not be surprising that one of the terminal repeats of the *PROZIP1*-LTR-RT has been integrated in another LTR-RT on chromosome 7 (Figure 1). TEs can be domesticated to the host’s own advantage (22). Here, TE proteins are often repurposed to play a role in the defence against pathogens, viruses and other TEs. However, our data suggest that *PROZIP1* emerged *de novo* from a terminal repeat sequence and not from a pre-existing coding region (34). Initially, it was thought that *de novo* gene evolution from non-coding regions was an unlikely event and that novel genes emerged through duplication and recombination of pre-existing coding regions (35). However, there has been a paradigm shift and it is currently accepted that *de novo* gene evolution from non-coding DNA is a continuous feature of eukaryotic genome evolution. For instance, in the recent divergence of *Oryza sativa* subsp. *japonica* it was inferred that on average 51.5 de novo genes per million years were generated and retained (36). *PROZIP1* has features that are more frequently found by *de novo* genes, as it contains one exon that encodes a small protein (137 amino acids) with high intrinsic structural disorder (27, 37, 38).

*PROZIP1* is *Zea* specific and likely evolved less than 728,000 years ago. Maize diverged from its close crop relative Sorghum (*Sorghum bicolor*) around 12 million years ago and encountered a round of allotetraploidization 5-12 million years ago (39, 40). Following allotetraploidization, the maize genome was further expanded through the proliferation of LTR-RTs, especially during the last three million years (41). Conceivably, *PROZIP1* is a product of this recent ongoing LTR-RT proliferation as numerous homologous LTR-RT of the *PROZIP1* LTR-RT could be found (Figure 2). In the last ∼728,000 years, the *PROZIP1* ancestor sequence encountered numerous mutations that gave rise to *PROZIP1* (Figure 2 - 3). However, in the period after the *PROZIP1* emergence, the last ∼128,000 years, very little mutations have been observed in *PROZIP1*. Of the 48 *PROZIP1* copies in 26 maize lines, 42 had an identical nucleotide sequence and 47 an identical protein sequence. Moreover, the *PROZIP1* nucleotide sequences in *Z. mays* subsp. *parviglumis* and *mexicana* were also identical to the allele most frequent in the maize population. This indicates that *PROZIP1* might subjected to selection pressure that conserves its sequence, especially as LTR-RTs are known to be prone to mutagenesis (41) (Figure 4).

**Figure 4.**
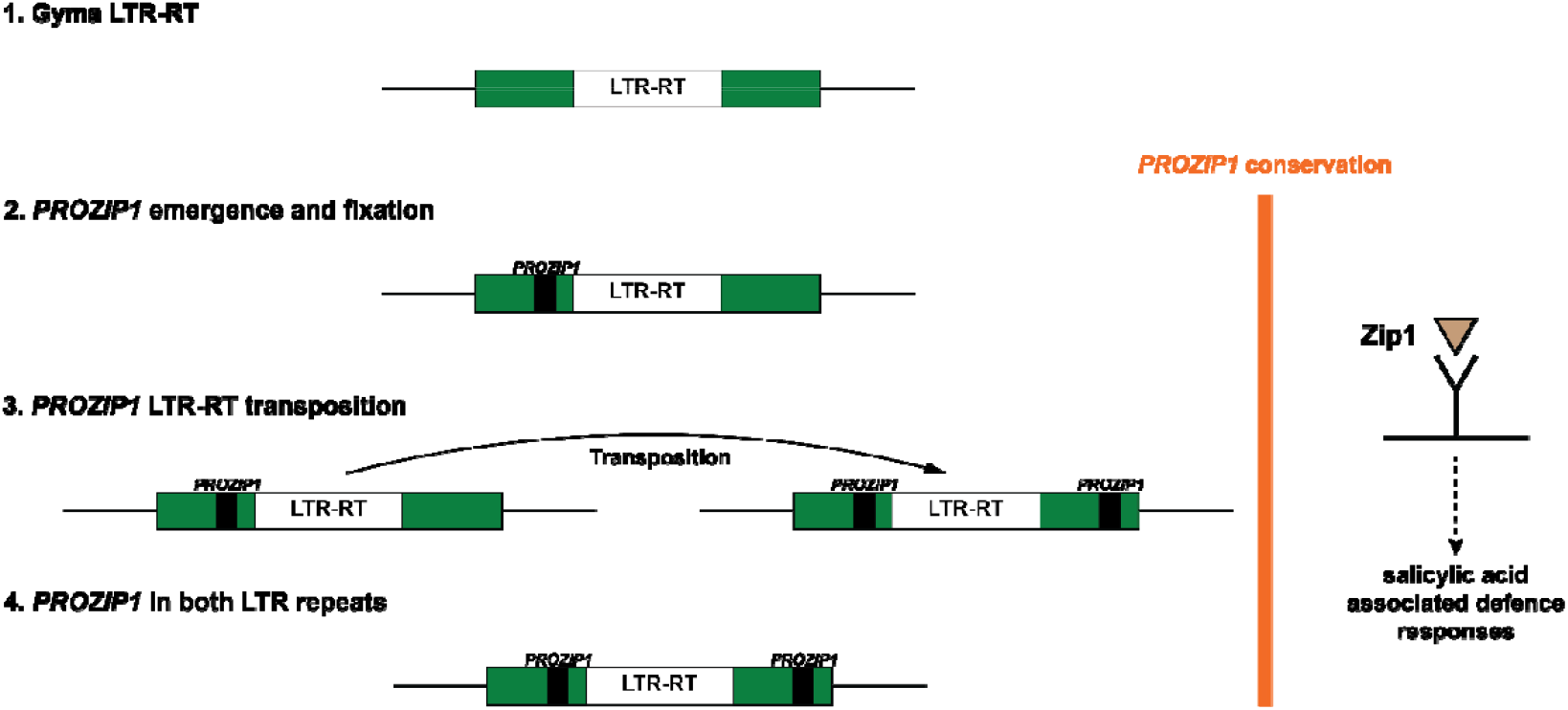
Model of the likely evolution of *PROZIP1*. (1) A Gyma long terminal repeat retrotransposons (LTR-RT) is subjected to mutagenesis leading to the continuous emergence of new open reading frames. (2) *PROZIP1* emerges and is fixated through its induction of salicylic acid associated defence responses. (3) The transposition of the LTR-RT with *PROZIP1* in the 5’ LTR leads to the presence of the LTR-RT at a different location with a *PROZIP1* copy in each of the two LTRs. (4) This LTR-RT is eventually fixated in most maize lines.

Zip1 elicits SA-dependent defence signalling in maize and may therefore be pivotal to fend off biotrophic plant pathogens (27). Sequence conservation of *PROZIP1* indicates the importance of this protein and its signalling role in the plant. However, the genome assemblies of P39 and Ky21 did not contain an open reading frame that can encode PROZIP1 (Table 1). The absence of *PROZIP1* in these maize lines might have consequences for their resistance against biological stressors. Indeed, maize line P39 and Ky21 have above average susceptibility against northern and southern leaf blight caused by the hemibiotroph *Setosphaeria turcica* and the necrotroph *Cochliobolus heterostrophus*, respectively (42, 43). This may indicate a role of Zip1 as facilitator of colonization of necrotrophic pathogens (27). However, this is certainly not the only factor determining susceptibility as there are NAM founder lines that contain *PROZIP1* and have a similar or higher susceptibility to leaf blight than P39 and Ky21 (42, 43).

TEs are generally more transcriptionally active under stressful conditions, which includes the attacks of biological agents (44). As any other genome sequence, retrotransposons are continuously subjected to mutagenesis that give rise to new open reading frame combinations that may occasionally provide an adaptive advantage (Figure 4). TEs may therefore be a good breeding ground for the emergence of new genes that encode small proteins that elicit particular responses in the plant. If these responses are elicited under conditions that provide an advantage to the plant, these open reading frames are more likely to be fixated. Plant genomic and proteomic data should be further scavenged for small proteins, which might have an impact on physiological processes. Moreover, we can study their genome locations and see if signalling proteins evolve more frequently *de novo* in association with transposons.

## Materials and Methods

### Genome assemblies, transposon annotation and classification

Assemblies of the 25 founder lines of the maize NAM population and a fifth assembly of the reference maize line B73 are publicly available, including their TE annotation (10). Homology and sequence identity between transposons were obtained through BLASTN (2.2.31+) (45). Transposable elements and homology between them were visualized using Biopython (v1.76) (46). Open reading frames in transposable elements and there encoding amino acid sequence of were determined with esl-translate (-l 50) as part of the Easel (v0.46) package. Functional domain within these amino acid sequences were determined with pfam_scan.pl (-e_seq 0.01) using the Pfam database version 35.0 (47).

### Ancestor sequence construction

The TE annotations of the NAM lines and B73 were used to identify the LTR-RTs (10). Homologous LTR-RTs were considered that have at least in 55% of their sequence more than 70% identity to the *PROZIP1* LTR-RT on chromosome 10 or, reversely, at least 55% of *PROZIP1* LTR-RT had more than 70% identity to homologous LTR-RTs. For this, BLASTN (2.2.31+) was used (45). Unique LTR-RTs were identified with Silix version 1.2.10-p1, using a minimum identity and overlap of 0.99 and 0.80 respectively (48). Only one representative of a sequence cluster was used. Homologous LTR-RTs were aligned using the software MAFFT (v7.464) (--auto --adjustdirection) (49). The phylogenetic tree and ancestor sequences were constructed using FastML (v3.11) (-- SubMatrix GTR --indelReconstruction BOTH) (50).

### SNode dating

Genome regions homologous to the terminal repeat of the *PROZIP1* LTR-RT were extracted from the maize teosinte *Zea mays* subsp. *mexicana* genome assemblies by using minimap2 2.17-r941 (51). To determine this sequence for *Zea mays* subsp. *parviglumis*, whole genome sequencing reads (SRR8181945) were mapped to the genome assembly of maize line Ky21(lacks *PROZIP1*, Table 1) with the addition of the *PROZIP1* LTR-RT terminal repeat that contains *PROZIP1* using Bowtie 2 (v2.3.5.1) (52, 53). Bcftools (v2.29.2) was then used to determine and incorporate the variants in order to construct a consensus sequence. Homologous sequences of the terminal repeat of the *PROZIP1* LTR-RT were aligned using MAFFT (v7.464) (--auto --adjustdirection) (49), trimmed with trimAl (v1.4.rev15) to remove sites with >20% gaps (54), and the phylogenetic tree was then constructed using RAxML (v8.2.11) with the nucleotide substitution model GTRGAMMA (55). The trimmed alignments were also used to determine the age of the different nodes using MrBayes (v3.2.7), with a HKY substitution model (nst = 2) and an inverse gamma prior on the clock rate (rates = invgamma) (56). Prior probability distributions were set for the branch lengths to be clock constrained uniform (brlenspr=clock:uniform) and the clock rate to have a normal distribution of 0.000001 ± 0.0000005 per site per 1000 years (clockratepr=normal(0.000001,0.0000005)). The node of the chromosome 10 allele and *Zea mays* subsp. *parviglumis* had an age assigned that was a normal distribution between 55,000 and 60,000 years that was used to calibrate the node dating.

## Acknowledgments

We thank Jeffrey Ross-Ibarra for helpful comments on the manuscript. This work has been supported by the Alexander von Humboldt Foundation, the German Research Foundation (DFG) via Project DO1421-/5-2 and the Cluster of Excellence on Plant Sciences (CEPLAS; Germany’s Excellence Strategy–EXC-2048/1 – Project ID: 390686111) and the University of Cologne.

## Supplementary Information

**Figure S1.**
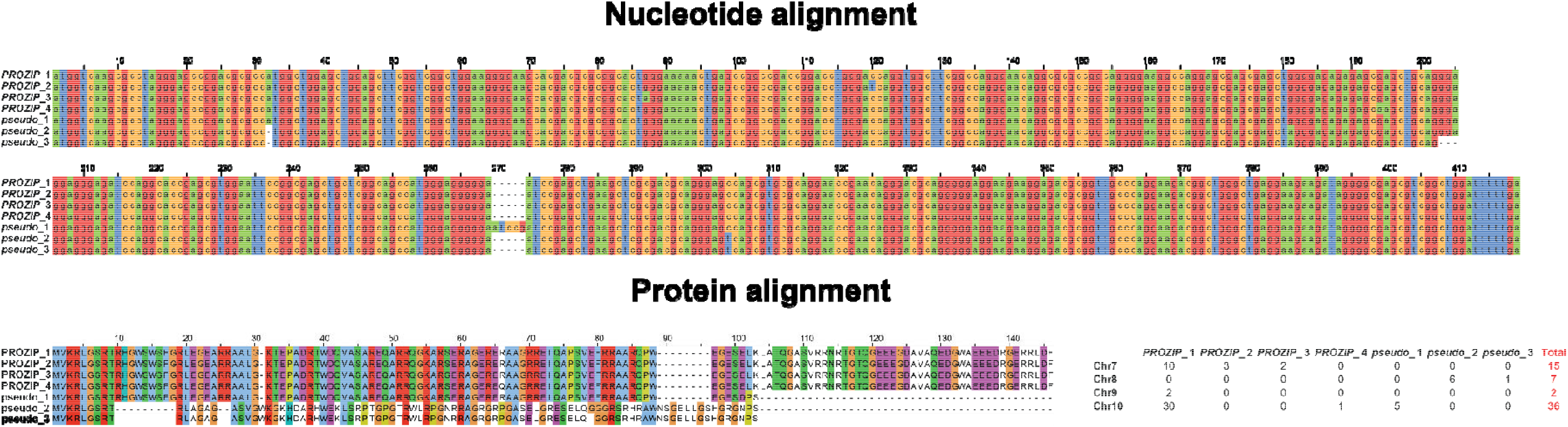
Alignment of *PROZIP1* and pseudo-*PROZIP1* alleles. The table in the right corner of the figure indicates the frequency of the different alleles.

**Figure S2.**
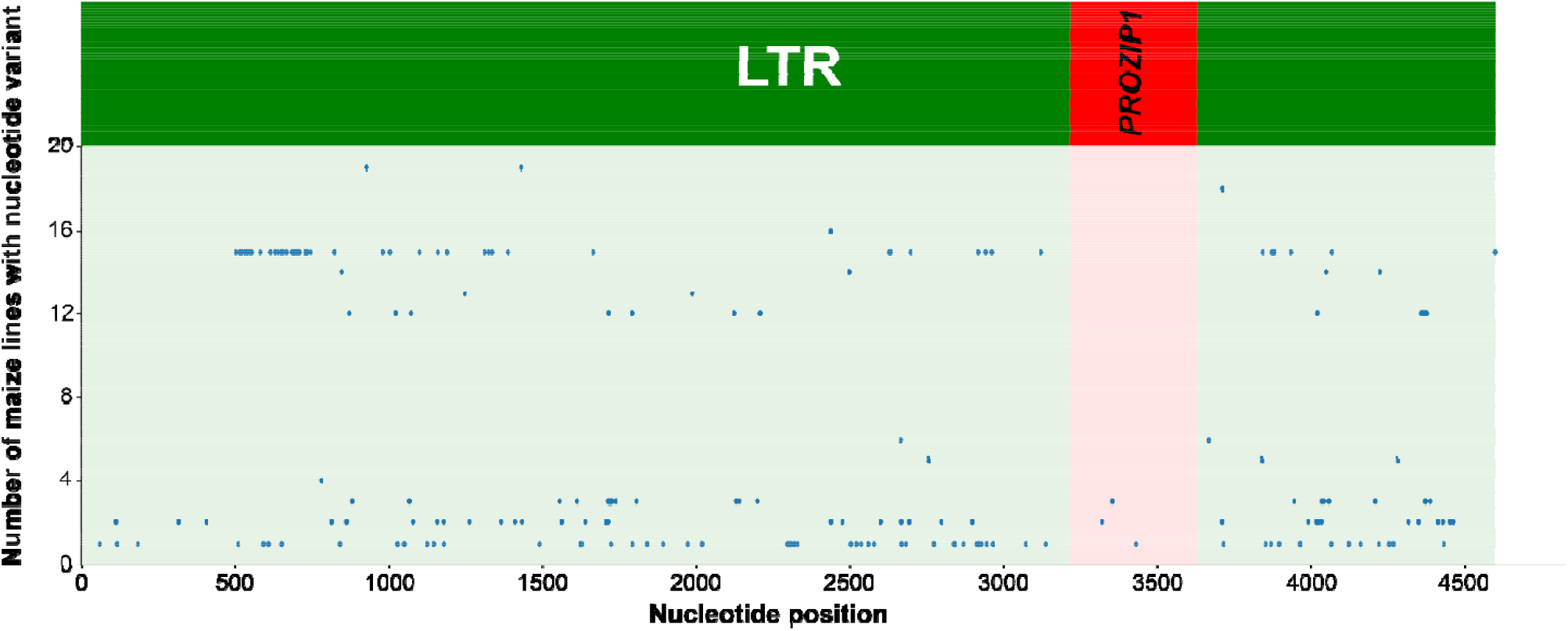
Nucleotide variants across the long terminal repeat where *PROZIP1* resides. Nucleotides are indicated that have variants in the NAM line and B73 population. A variant can be a nucleotide polymorphism or an indel (insertion or deletion).

